# Estimating barriers to gene flow from distorted isolation by distance patterns

**DOI:** 10.1101/205484

**Authors:** Harald Ringbauer, Alexander Kolesnikov, David Field, Nicholas H. Barton

## Abstract

In continuous populations with local migration, nearby pairs of individuals have on average more similar genotypes than geographically well separated pairs. A barrier to gene flow distorts this classical pattern of isolation by distance. Genetic similarity is decreased for sample pairs on different sides of the barrier and increased for pairs on the same side near the barrier. Here, we introduce an inference scheme that utilizes this signal to detect and estimate the strength of a linear barrier to gene flow in two-dimensions. We use a diffusion approximation to model the effects of a barrier on the geographical spread of ancestry backwards in time. This approach allows us to calculate the chance of recent coalescence and probability of identity by descent. We introduce an inference scheme that fits these theoretical results to the geographical covariance structure of bialleleic genetic markers. It can estimate the strength of the barrier as well as several demographic parameters. We investigate the power of our inference scheme to detect barriers by applying it to a wide range of simulated data. We also showcase an example application to a *Antirrhinum majus* (snapdragon) flower color hybrid zone, where we do not detect any signal of a strong genome wide barrier to gene flow.

---

Many populations are distributed across geographically extended habitats that are sometimes interrupted by barriers to gene flow. They can arise due to physical obstacles that reduce migration, but can also be caused by genetic incompatibilities, which reduce gene flow across a hybrid zone (Barton 1979). Barriers can prevent locally adapted populations from being swamped by dispersal, and they can facilitate divergence, ultimately leading to speciation. Therefore, they play a central role not only in conservation, but also in evolutionary biology and ecology. As direct observations of individual movement and reproduction are time-consuming and expensive, and moreover can only give a snapshot in time, there is much interest in indirect methods that infer such barriers from observed geographic genetic structure.

Such methods to detect barriers from genetic data can be grouped into two distinct approaches (Guillot *et al.* 2009): Clustering methods, that detect geographic genetic discontinuities between populations by grouping individuals into population units based on genetic similarity (Falush *et al.* 2003; Guillot *et al.* 2005; Dupanloup *et al.* 2002), and edge detection methods, that identify areas of sharp genetic change (Womble 1951; Cercueil *et al.* 2007; Manni *et al.* 2004). None of these approaches is directly linked to any spatial population genetic model. They can therefore infer the existence of a barrier, but cannot give meaningful and biologically interpretable estimates of its strength. In addition, these approaches are often confounded by isolation by distance patterns (Safner *et al.* 2011; Meirmans 2012), whereby individuals nearby are more similar than distant individuals (Wright 1943) due to recent co-ancestry. While the description of this effect has a long history in theoretical population genetics of homogeneous populations (Malécot 1948; Slatkin 1993; Rousset 1997; Hardy and Vekemans 1999; Barton *et al.* 2002), it has not been included into a practically applicable method to estimate the strength of a barrier to gene flow.

Here, we fill this gap, and introduce a method that infers the strength of a barrier in a two dimensional population by fitting a population genetic model. Our method utilizes the fact that a barrier to gene flow distorts classical isolation by distance patterns (Fig. 1). Based on theoretical work of (Nagylaki 1988), Barton (2008) constructed a theoretical framework. He showed that in two spatial dimensions, where fluctuations of allele frequencies are more localized than in one dimension, these effects of a barrier on allele frequency fluctuations can be significant already for intermediate barrier strengths. This signal therefore holds big potential for demographic inference. The derivation of Barton (2008) also shows that the effect of a barrier depends primarily on short-lived, localized fluctuations. In general, isolation by distance patterns equilibrate relatively quickly and depend mostly on recent demography (Aguillon *et al.* 2017; Barton *et al.* 2013). Therefore, an inference scheme based on distorted isolation by distance patterns infers contemporary barriers to gene flow, and should be robust to confounding effects of ancestral structure.

**Figure 1.**
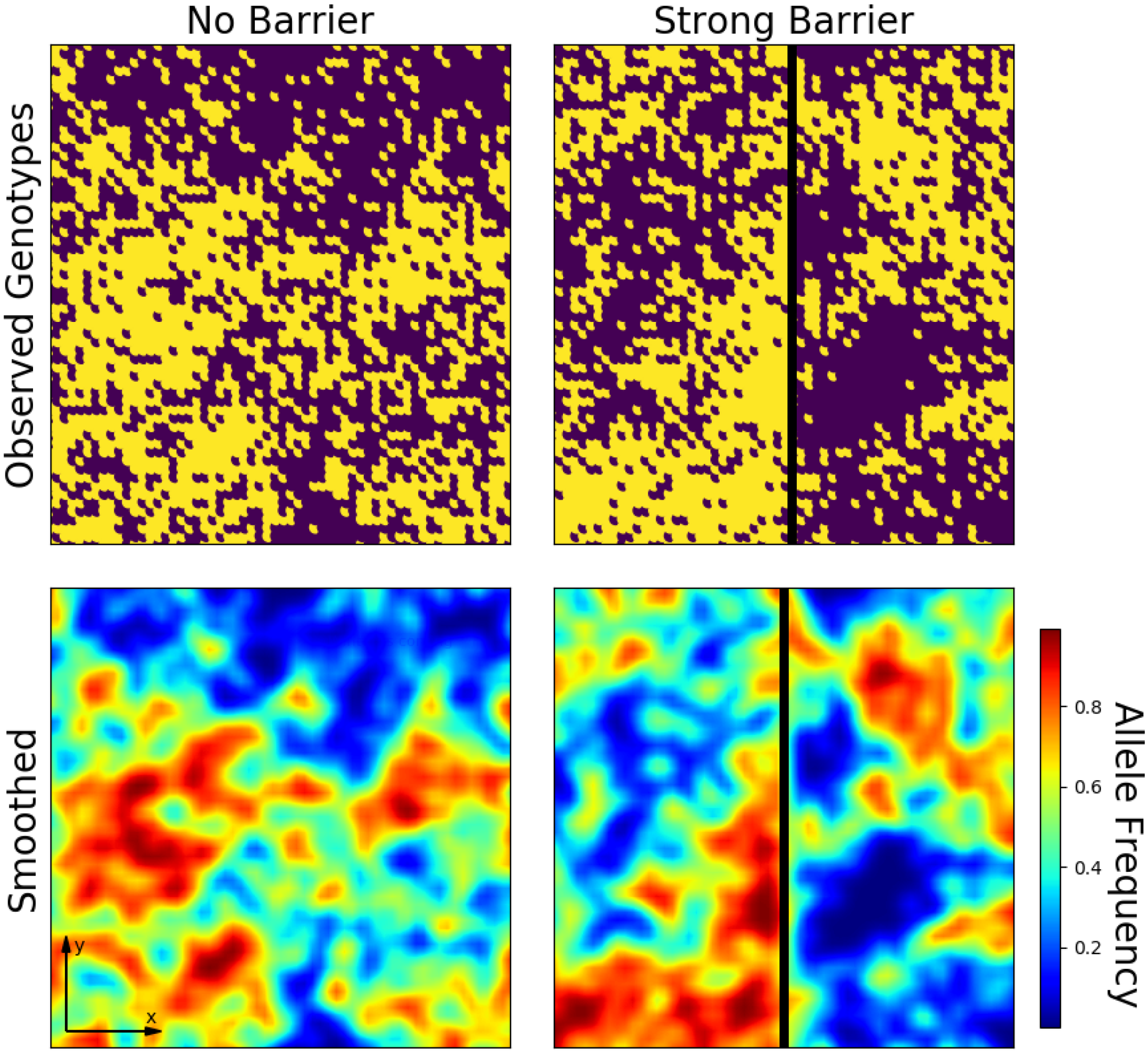
Geographic fluctuations of allele frequencies. We simulated individual discrete genotype data for one locus in a two-dimensional habitat in absence of a barrier to gene flow (left) and in presence of a strong barrier (right). We used the backward simulation scheme outlined below with a axial variance of dispersal *σ*^2^ = 1, one individual per deme (*D*_*e*_ = 1), a long distance migration rate *μ* = 0.001 and a complete barrier *κ* = 0. Top: Discrete genotype data. Bottom: Allele frequencies are smoothed with a Gaussian Kernel for better visualization. This figure demonstrates the underlying idea of our method: A barrier distorts random geographic fluctuations of allele frequencies. The strength of these local fluctuations increases on average next to the barrier, and there is less correlation across the barrier. This signal can be used to infer the strength of the barrier.

Here, we first expand previous theoretical results that describe the effect of a barrier on classical isolation by distance patterns (Nagylaki 1988; Barton 2008). We introduce a model where ancestry diffuses backwards in time and is partially reflected by a barrier. This allows us to numerically calculate the probability of recent co-ancestry, which can then be fitted to genetic data. As single nucleotide polymorphism (SNP) datasets are currently widely used, we develop and implement ways to fit such biallelic genetic markers.

We test our inference scheme on synthetic data simulated under an explicit population genetics model and investigate how it is affected by confounding factors, for instance in a scenario of secondary contact. We also show a practical application, in which we apply our inference scheme to a hybrid zone population of *Antirrhinum majus,* in which a sharp transition in flower color and a causal flower color gene occurs (Whibley *et al.* 2006). We apply our method to test whether there is also a genome wide barrier to contemporary gene flow. To this end, we analyze a dataset of 12389 individuals and 60 suitable SNP markers.

## Materials and Methods

We first outline the model underlying our inference scheme and discuss its assumptions, and then describe how our method fits this model to observed genotype data. In brief, we use a diffusion approximation for the spread of ancestry to calculate the probability of recent identity by descent between pairs of samples. We then fit our model to data by finding the demographic parameters that maximize the fit of observed homozygosity between all sample pairs.

### Model

No model can capture all complexities of the real demographic history of a population. Therefore, the aim is not to have a mathematically exact model, but one that robustly captures general patterns of spatial fluctuations of allele frequencies. We use a model of a two-dimensional continuous habitat that is interrupted by a barrier and assume that the demographic parameters are the same on both sides. In short, we calculate the chance of pairwise coalescence before a long distance migration or mutation event. We use a diffusion model to trace lineages backwards in time, and assume that rare long distance migration events, which counteract the build up of local allele frequency fluctuations, occur at a constant rate. To calculate the equilibrium identity by descent pattern, i.e. the probability that two lineages coalesce before a rare long distance migration or mutation event happen, we first derive the coalescence probability at specific times *t* in the past and then integrate over *t*.

Our model is closely related to previous theoretical treatments of allelic identiy by state in presence of a barrier to gene flow (Nagylaki 1988; Barton 2008). For a population occupying a linear habitat, Nagylaki (1988) derived continuous equations for identity by state by taking the limit of a model of linear demes that exchange migrants (the so called stepping stone model). Barton (2008) expanded this by solving an analogous equation for two dimensional populations. His formulas are given as a numerical Fourier transform that diverges for nearby individuals. These equations for two dimensional populations are formally problematic, as they were not obtained by rescaling (which is impossible in two spatial dimensions), but Barton (2008) demonstrated that the solution is in close agreement with the solutions from a discrete stepping stone model for all but very close distances.

Here, we base our inference scheme on a different approach. We use a diffusion approximation to describe the spread of ancestry backwards in time. While the results are formally equivalent, we found our approach to be computationally more robust, and, most importantly, more efficient. This approach allows us to apply our method to sample sizes of hundreds to thousands of individuals, and dozens to hundreds of loci.

#### Diffusion approximation

We model the spread of ancestry using a geographic diffusion approximation, which has a long history in population genetics (Fisher 1937; Wright 1943; Malécot 1948; Nagylaki 1978). Tracing a lineage at one locus backwards in time, the total spatial movement of the lineage is the sum of many independent migration events. If these events are sufficiently uncorrelated, the overall spread of ancestry can be approximated as a random walk process. This approximation will be accurate as long as rare large scale events do not significantly influence the movement of ancestral lineages. Therefore, the diffusion approximation is expected to be accurate on recent to intermediate time scales, on which large scale events such as colonizations often play only a minor role.

In the absence of a barrier, the process is a free diffusion, and the probability density function (PDF) of finding an ancestor at position *y* at time *t* along a given axis is given by a Gaussian probability density function arount the current position *x* and variance *σ*^2^*t* that increases linearly backwards in time:

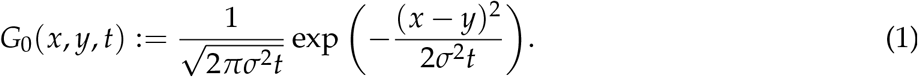

The dispersal rate *σ* describes the speed of the spread of ancestry. In case of a homogeneous density of individuals across the landscape, the backward dispersal probability density equals the probability density of lineages moving forward in time. The dispersal rate *σ*^2^ can be interpreted as the axial variance of the one generational dispersal kernel then, if time is measured in generations (Rousset 1997). The diffusion approximation is fully determined by the equation:

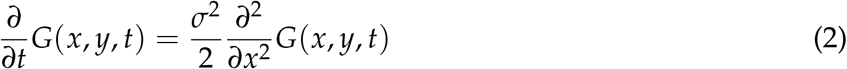

with initial condition:

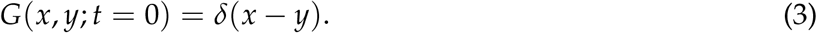

#### Partial barrier to gene flow

We model a barrier as a partially permeable barrier to diffusion of ancestry. For a barrier at *x* = 0, the following interface boundary conditions have to be supplied (Grebenkov *et al.* 2014):

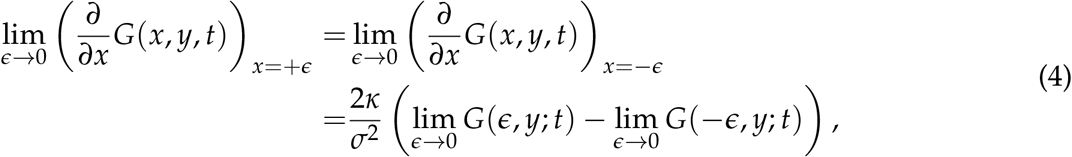

The first line describes the constancy of the flux across the barrier. For *κ* = 0 there is no flux across the barrier, and the barrier is infinitely strong. On the other hand, the case *κ* = *∞* implies the continuity of the probability density across the barrier and the solution reduces to free diffusion. Comparing to differential equation (54) of Nagylaki (1988), which is derived by rescaling a stepping stone model, gives an intuitive interpretation of *κ*: This parameter corresponds to the fraction of successful migrants across a barrier if demes are spaced one dispersal unit apart. The quotient 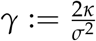 corresponds to an equivalent factor in formula (54) of Nagylaki (1978). It is also the inverse of the barrier strength parameter *B* defined by Barton and Bengtsson (1986), which has dimension of distance.

Equations 2, 3 and 4 allow for an analytic solution for the probability density with a barrier (Grebenkov *et al.* 2014):

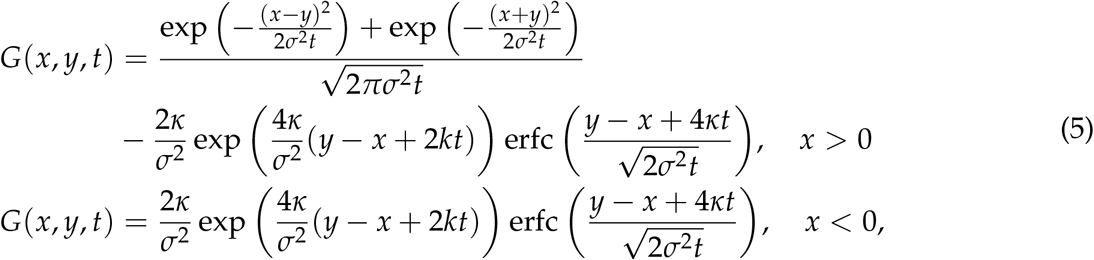

where erfc(*z*) denotes the complementary error function. These expressions are valid for *y* > 0, and their extension to *y* < 0 is straightforward by the symmetry *x, y* → –*x*, –*y*. In Fig. 2, these formulas are compared to random walk simulations. The PDF converges to the Gaussian of free Brownian motion for *κ* → 0. For a two-dimensional diffusion process with a linear barrier at *x* = 0, the full solution is given by multiplying the one-dimensional density functions Eq. 2 for movement parallel to the barrier and Eq. 5 for movement normal to the barrier.

**Figure 2.**
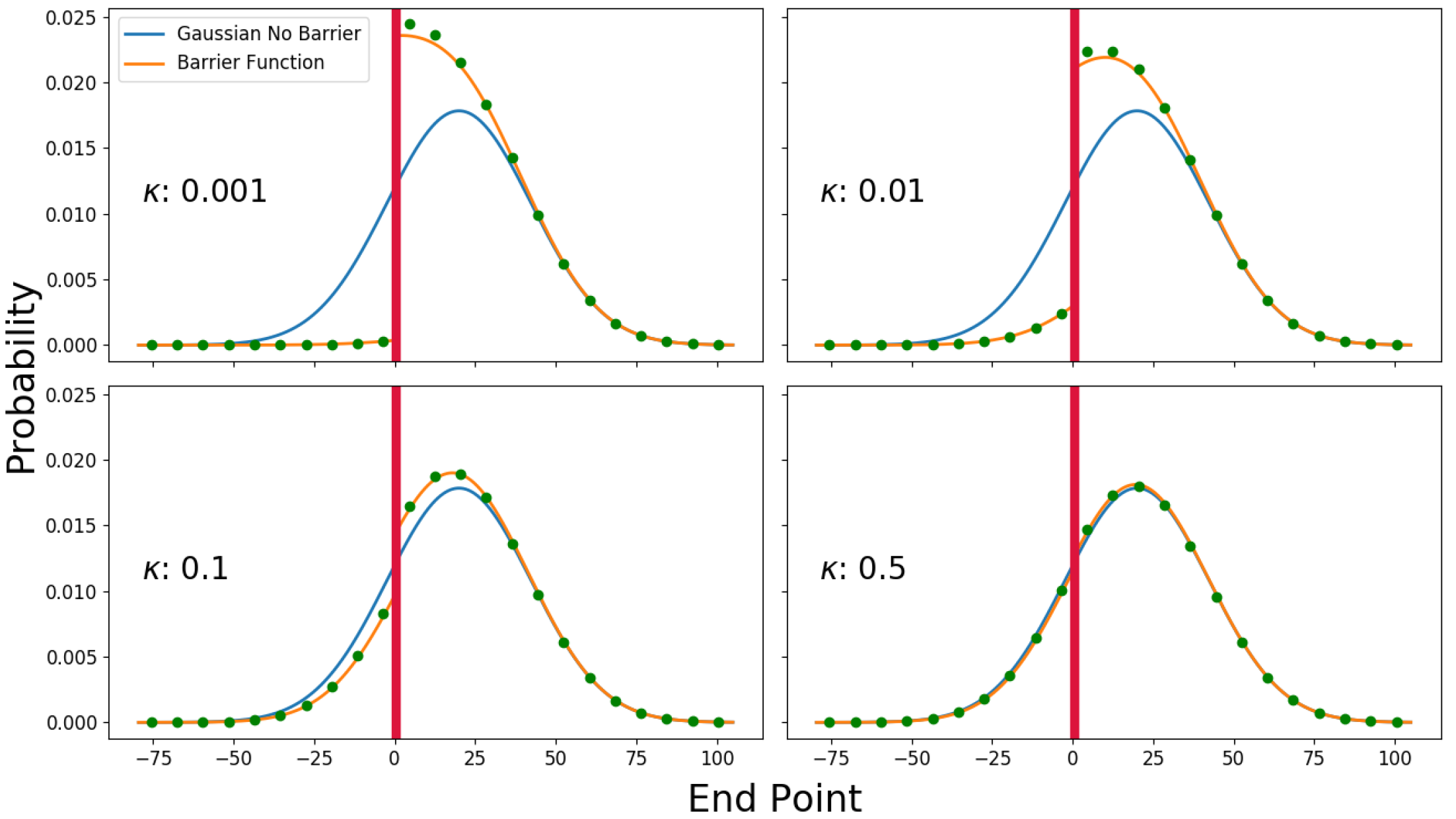
Comparison of analytical diffusion formulas (Eq. 4) with discrete random walk simulations for *t* = 500 in the past. We simulated a one-dimensional random walk on an array of linear discrete nodes and a barrier at *x* = 0. Every generation, a random step to one side is made (which implies that *σ* = 1). If a movement would be across the barrier, it is realized with probability *κ*. For each of four barrier strengths, we simulated 10^6^ replicates starting at *x* = 20. The blue line depicts the corresponding Gaussian probability density of free diffusion in absence of a barrier.

#### Pairwise coalescent probabilities

We use the diffusion approximation to model the distribution of coalescence times for pairs of individuals. We approximate the chance of co-ancestry originating in a small time interval *dt* around time *t* ago as the product of the probability of coming close and a rate of local coalescence 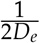 (Ringbauer *et al.* 2017). The local density *D*_*e*_ describes a rate with which nearby lineages coalesce (Wright 1943). In a stepping stone model, *D*_*e*_ corresponds to the number of diploid individual per deme (Barton *et al.* 2002). This approximation ignores that lineages do not move apart again once they have coalesced. An equivalent simplification is made by Barton (2008), and it is accurate as long as coalescence is sufficiently rare (Wilkins 2004). Since the dispersal process is symmetrical in time in our model of constant population density, the chance that two lineages at current position *x* and *y* are close at time *t* back equals the probability density that a single lineage moves from *x* to *y* in time 2*t*. Formally, the probability density *ψ*(*x, y, t*) of coalescence *T*_*c*_ at time *t* in the past is approximated by

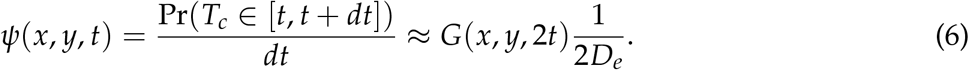

#### Identity by descent

We define identity by descent *F* of two samples at 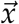 and 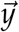 as the chance that two lineages coalesce before a long distance migration or, equivalently (but unlikely for single nucleotide polymorphisms), a mutation event occurs along one of the lineages. This definition is closely related to the widely used fixation index *F*_*ST*_, and both definitions agree in the limiting case of an infinite population (see Rousset (2002) for a review). If one assumes that mutation or long-distance migration occur at a constant rate *μ*, it is straightforward to calculate the probability that two lineages coalesce before a long distance event:

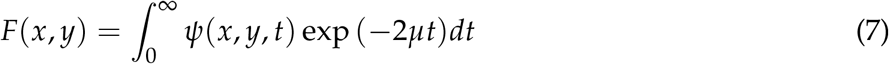

In the absence of a barrier, the probability of identity by descent varies only with the Euclidean distance between two individuals. In this special case, the integral in Eq. 7 has an analytical solution, the classical Wright-Malecot formula (Barton *et al.* 2002, 2013):

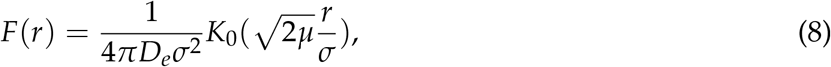

where *K*_0_ is the modified Bessel function of the second kind of degree zero. A caveat of this analytical solution is that *K*_0_ diverges logarithmically as *r* → 0 (Barton *et al.* 2002). Similarly, the integral in Eq. 7 diverges for nearby individuals. As for the Wright-Malecot formula, this is caused by a behavior of the diffusion approximation for short timescales, as in this model the chance of two lineages being close diverges as 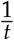 for *t* → 0. An obvious solution to circumvent this problem is to start integration at time *t*_0_ > 0. Here, we choose one generation time, a biologically plausible value. We could not find an analytical solution, but Eq. 7 can be numerically integrated.

Our results show that if a barrier to gene flow is present, identity is decreased across the barrier, and increased for points on the same side of the barrier (Fig. 3). Interestingly, the increase of *F* for a pair of points 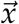 and 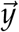 on the same side of the barrier equals the decrease of identity between points 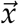 and 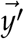, where 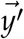 is the point 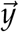 reflected across the barrier. This symmetry originates from a reflection principle of the underlying random walk model, as lineages that do not cross the barrier behave as if they were reflected. This symmetry already occurs in the barrier point density function (Eq. 5). It implies that for a complete barrier identity by descent can increase to at most twice of the value in absence of a barrier, as observed in the equivalent case of a range boundary (Wilkins 2004).

**Figure 3.**
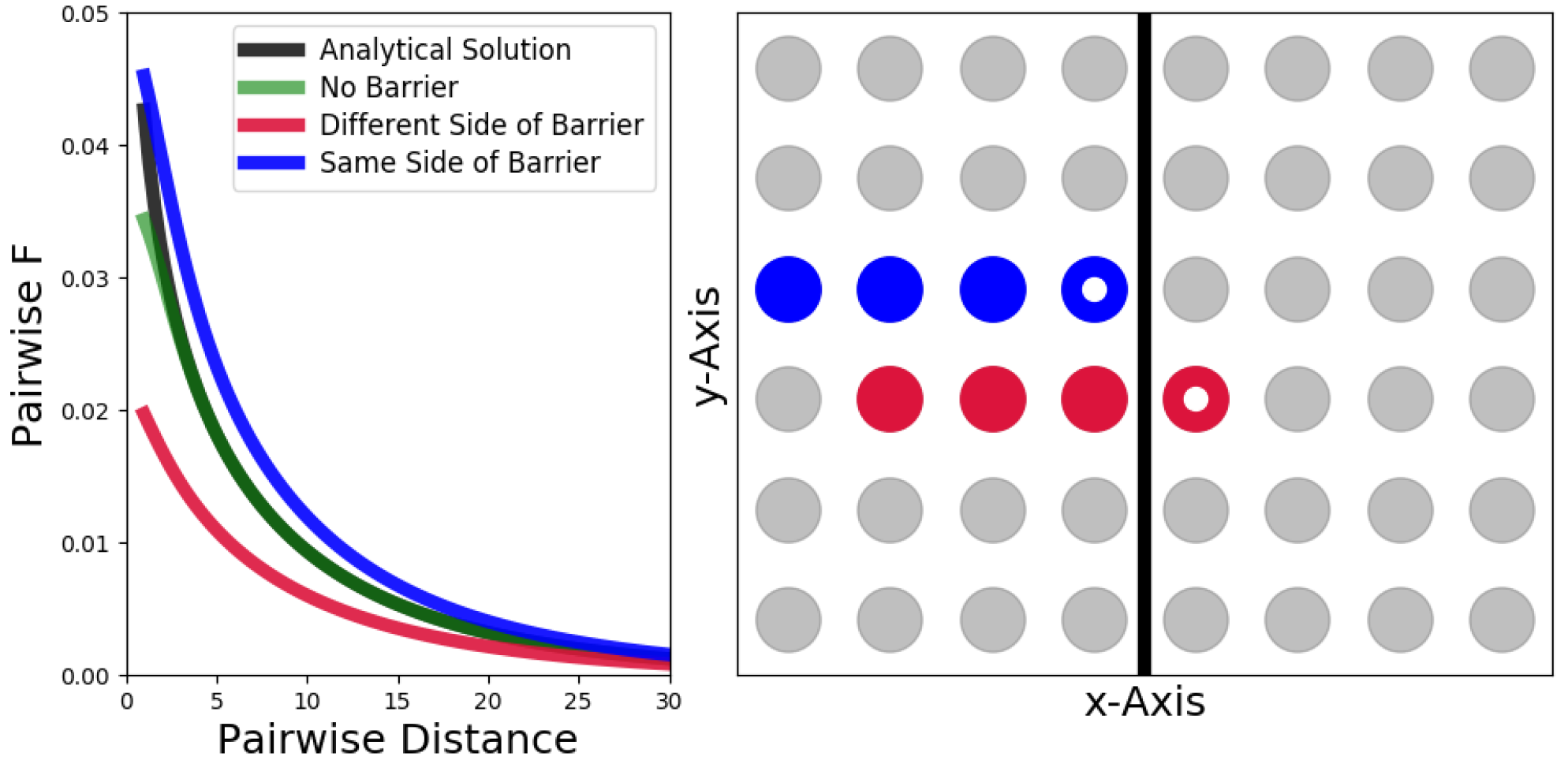
Decay of identity by descent *F* in presence of a strong barrier to gene flow and moderate neighborhood size (*κ* = 0.1, *σ* = 1, *D*_*e*_ = 5 and *μ* = 0.003). Individuals differ in their *x*-coordinate, and we calculate *F* for an individual at *x* = 1 and an individual at x=–1 with other individuals to the left of the barrier (right). The analytical solution in the absence of a barrier is calculated via the Wright-Malecot formula (Eq. 8), the others are integrated numerically via Eq. 7.

#### Rescaling

Not all parameters in Eq. 7 are independent. Consequently, they cannot be estimated separately, as in absence of a barrier (Rousset 1997; Barton *et al.* 2013). Therefore, we replace the four demographic parameters 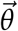: *D*_*e*_, *κ*, *μ*, *σ* in equation Eq. 6 with three compound parameters 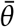: Neighborhood size Nbh:= 4*D*_*e*_*σ*^2^*π* (a classical parameter that goes back to Wright (1943)), a scaled barrier parameter 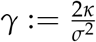 (corresponding to the inverse of Barton’s *B*), and a scaled long-distance migration rate 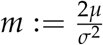 (Appendix).

### Fitting the Model to Data

A typical dataset consists of genotypes 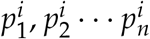 for a marker *i* and individuals at positions 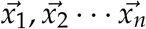. To infer the underlying demographic parameters 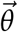 from observed data, we have to develop a way to fit our model to such data.

In principle, it is straightforward to transform the probability of identity by descent as calculated by our model into expected allele frequency covariances. For two samples *k* and *l*, the expected covariance between their genotypes at marker *i* around a population mean allele frequency *p̄*_*i*_ is given by:

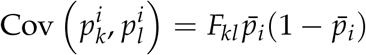

A serious problem with this approach is that often mean allele frequencies have to be estimated from the data, but estimating the means of allele frequencies for many markers would lead to severe over-fitting. In order to circumvent this caveat, we developed and tried different methods to fit identity by descent to genotype data without directly estimating all mean allele frequencies. We included one approach that models individual genotypes as binomial draws from latent allele frequencies modeled by a Gaussian random field (Supp. Text 1).

#### Fitting pairwise homozygosity

Of all tested fitting methods a relatively simple approach that fits the fraction of pairwise homozygosity (defined as fraction of identical genotypes) has least bias and sampling variation (Supp. Text 1). Throughout this work, we use this method for data analysis. In the following we give a brief outline of how we calculate expected homozygosity and how we fit it to data.

The observed average homozygosity *h*_*kl*_ for a pair of individuals *k* and *l* with genotypes 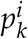 and 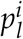 at markers *i* = 1,2,…, *n* can be straightforwardly calculated from the data:

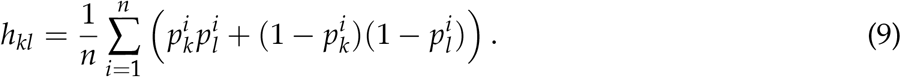

Our model predicts the pairwise chance of identity by descent *F*, and these probabilities can be used to calculate the expected values of average homozygosities *E*(*h*_*kl*_). Denoting the probability of identity by descent between a pair of samples *k* and *l* by *F*_*kl*_ and the mean allele frequency of a single marker *i* by *p̄*_*i*_, the expected pairwise homozygosity at this marker is given by:

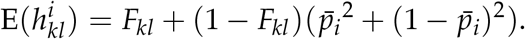

The first term gives the probability of having the same genotype due to identity of descent and the second describes the probability of having the same genotype by chance. To avoid estimating all mean allele frequencies, we sum over all markers to get the expected fraction of pairwise identical genotypes:

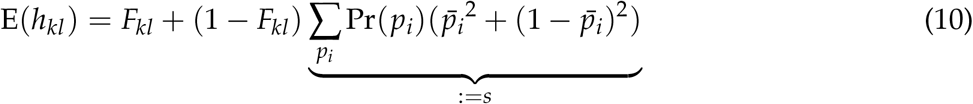

Instead of fitting all unknown allele frequencies, now only one additional compound parameter *s* has to be fit in addition to the demographic parameters *θ*. We tried fitting this formula to observed data with a composite likelihood approach. However, we found that minimizing the sum of all squared deviations between the expected and observed pairwise homozygosities

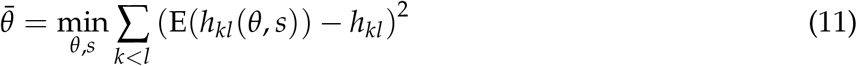

gives almost equivalent results, while being additionally much faster (Supp. Text 1).

Fitting pairwise homozygosities can be easily extended to deme data, where nearby individuals have been binned, by plugging deme allele frequencies into Eq. 9.

#### Estimation uncertainties

To learn about estimation uncertainty, we bootstrap over genetic markers. Unlinked markers contain almost independent information because their spatial movements are typically correlated only on very short timescales. Therefore, resampling loci at random is expected to yield accurate empirical confidence intervals.

### Implementation

In brief, our inference scheme does the following three computational steps for a given set of demographic parameters 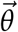:

1. Calculate pairwise *F* for all pairs of samples (integral Eq. 7).
2. Use these pairwise *F* to calculate the expected pairwise homozygosity for all pairs of samples (Eq. 10).
3. Calculate the sum of squared differences between the expected and observed pairwise homozygosities (Eq. 11).

Our program then finds the parameters 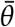 that minimize this function by using the Levenberg-Marquardt algorithm, as implemented in the Python package *Scipy*.

We implemented the described simulation and inference methods mostly in Python. To speed up calculations we parallelized the calculations for pairwise *F*, so that they can be run simultaneously on different CPUs. The evaluation of the integrand Eq. 7 is a computational bottleneck. We implemented this calculation in C, to make use of the superior speed of a compiled language.

The inference scheme has to compute the expected identity by state for every pair of samples. It therefore scales quadratically with the number of individuals, as there are 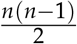 such pairs. It has a runtime of several hours for a individual/deme number of 1000 when run on a single standard desktop CPU. In order to produce a sufficient number of replicates and bootstraps, we utilized a scientific computer cluster at IST Austria. To speed up runtime, individuals can be grouped into demes. If clustering is done on small scales for bins at most a few *σ* in diameter, it does not significantly affect the estimation scheme (Supp. Text 1). In our results, we capped *γ* to a maximum value of 1, as for *γ* > 1 the effect of a barrier becomes negligible (Fig. 2).

### Simulations

We extensively tested our inference scheme on simulated data sets. We used a stepping stone model with *D*_*e*_ individuals per deme, and we traced ancestry backwards in time. Every generation each individual picks an ancestor at random with probabilities given by a dispersal kernel. We mostly used the heavy-tailed Laplace distribution, but due to the rapid convergence to the continuous diffusion approximation (Fig. 2), the specific choice has no significant impact. If two lineages happen to pick the same ancestor, they coalesce into one ancestral lineage. We simulate long distance migration events to occur at a constant rate, and they make the corresponding lineage pick an allele at random from the population mean allele frequency *p̄*. In order to model the effects of a barrier, we follow Nagylaki (1988) and realize migration events across the barrier only with relative probability *κ*, the barrier strength parameter. For constant deme sizes, this backwards models is equivalent to a forward model in which a large number of gametes are dispersing with the same dispersal kernel (Nagylaki 1988).

After a preset maximum number of generations, every lineage picks an allele at random according to the mean allele frequency *p̄*. Different, unlinked loci were simulated as independent runs. We picked mean allele frequencies at random according to a predetermined distribution, usually Gaussian with standard deviation *σ*(*p̄*) around an overall mean of 0.5. We also investigated how robust our model is to scenarios of secondary contact (Fig. 3). To this end, lineages were assigned an allele with probability *p̄*_*l*_ or *p̄*_*r*_, according to their location at time of secondary contact.

We stress that the data is simulated under a process very similar to our model. For real data, there could be other deviations from the model which might further reduce the power of the inference scheme. Therefore, our results should be seen as limits for the inference scheme in case of ideal data.

### Anirrhinum majus Data

To show the practical utility of our inference scheme, we applied it to data from an *Antirrhinum majus* hybrid zone. This hybrid zone is located in a valley in the Eastern Pyrenees. It shows a geographically narrow transition between two flower color morphs, in which a range of hybrid flower color phenotypes occur. This transition is mainly determined by three major flower color genotypes that regulate the intensity and patterning of flower color (Whibley *et al.* 2006). We applied our method to a dataset of 12389 plants collected between 2009 and 2014, which were genotyped for 112 SNP markers.

To satisfy the assumptions of our model as good as possible, we filtered markers based on four quality criteria: minor allele frequency, large scale geographic correlation, linkage disequilibrium and deviations from local Hardy Weinberg equilibrium. SNP design and filtering are explained in detail in Supp. Text 3. After this data cleaning step, we were left with 60 unlinked polymorphic SNPs, that were spaced throughout most of the genome (Supp. Text 3).

### Data Availability

The source code for the implementation of our inference scheme is freely accessible at the Github repository https://github.com/hringbauer/BarrierInfer.git

The *Antirrhinum majus* dataset is a subset of samples collected from 2009 to 2014 with the long term goal to build a pedigree. The details of this dataset and data filtering are described in Supp. Text 3.

## Results

### Inference on Simulated Data

We investigated the overall capability of this method to estimate barrier strengths and the accuracy of empirical bootstrap uncertainty estimates. Our tests show that the inference scheme can reliably recover barrier strengths as well as demographic parameters (Fig. 5 and Fig. 6). Estimates of the neighborhood size are robust, but show a slight upward bias. These slight biases are likely due to the fact that a continuous model is used to fit to discrete simulations. Estimates of the long distance migration rate *m* are more variable, but are not significantly biased. In all cases, the range of bootstrap estimates mostly overlaps with the true value to the expected degree. This result indicates that bootstrapping allows one to get accurate uncertainty estimates (Fig. 6).

The stronger the barrier, the more strongly it effects allele frequency fluctuations. Our results indicate that the inference scheme has higher power to infer strong barriers (*γ* < 0.1), whereas weaker barriers cannot be inferred reliably (Fig. 5). The exact power of the method will depend on a combination of several factors, in particular the strength and the geographic extent of isolation by distance patterns, as well as the geographic sampling scheme. As a barrier mostly affects fluctuations near it, generally a high sampling density next to the putative barrier is preferable. Our power simulations for a specific, realistic scenario indicate that at least a few dozen markers and several hundred individuals are required for robust inference of strong barriers (Supp. Text 1).

#### Secondary contact

Barriers to gene flow sometimes coincide with areas of secondary contact, for instance in secondary hybrid zones (Barton and Hewitt 1985). Allele frequencies might have diverged before this contact, and present day allele frequency differences are not caused by the presence of a barrier. However, these clines resembles the effect of a barrier, and might be mistakenly inferred as such. One salient way to deal with this problem is to remove markers that show large scale geographic structure of allele frequency. One can base inference on a subset of markers that have similar mean allele frequency across the whole population range and only display fluctuations on small geographical scales, which equilibrate quickly (Barton *et al.* 2013). We tested this approach on simulated data. When applying the inference scheme to a simulated scenario of secondary contact with divergent allele frequencies, it wrongly infers a barrier in case of no filtering (Fig. 7). However, when using the subset of loci that show no large scale correlation with geography, the false positive signal decreases. Moreover, filtering out loci with large scale structure does not remove the signal in case of a true barrier since secondary contact (Fig. 7). However, if sampling is only done on small spatial scales, such filtering could become problematic, as one might remove signal from local fluctuations as well. Therefore we advise to always check that the sampling area is bigger than the spatial scale of isolation by distance patterns before any markers are removed.

**Figure 4.**
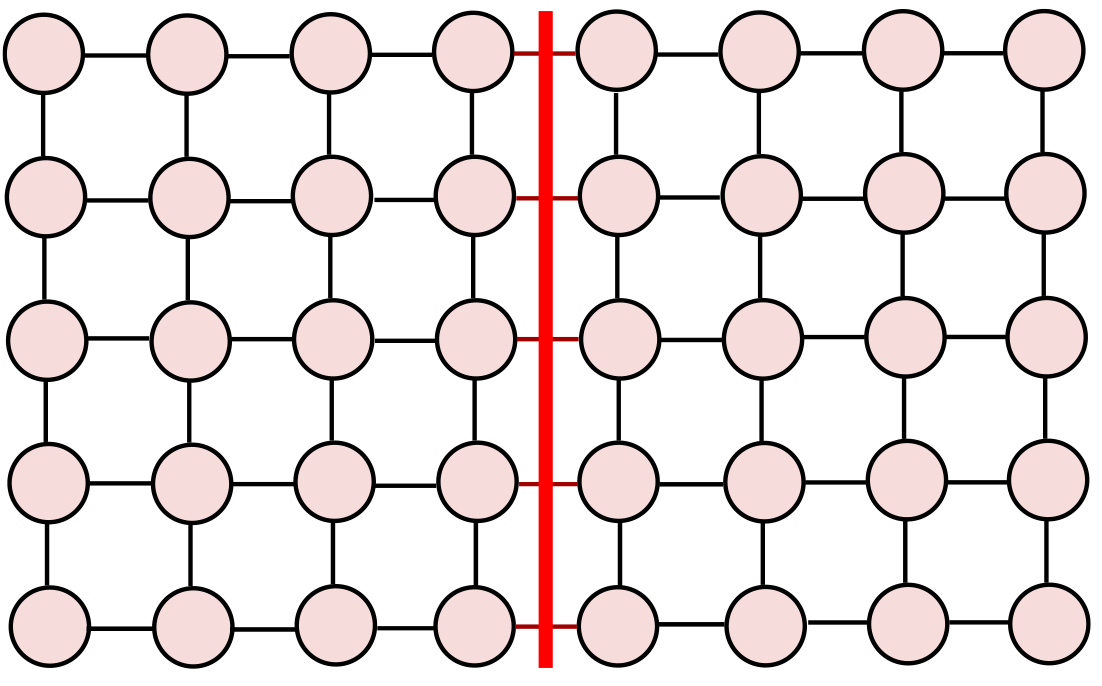
Model used to generate synthetic data sets. Ancestry is traced back on a twodimensional grid of demes in discrete time steps. An ancestral deme is randomly picked according to a dispersal kernel, and then an ancestor is chosen at random from within this deme. If lineages fall on the same ancestor, they coalesce into a single lineage. For each unique lineage and every step, there is constant chance that a long distance migration event occurs. In this case, the lineage and all corresponding individuals pick the same allele randomly drawn from a mean allele frequency.

**Figure 5.**
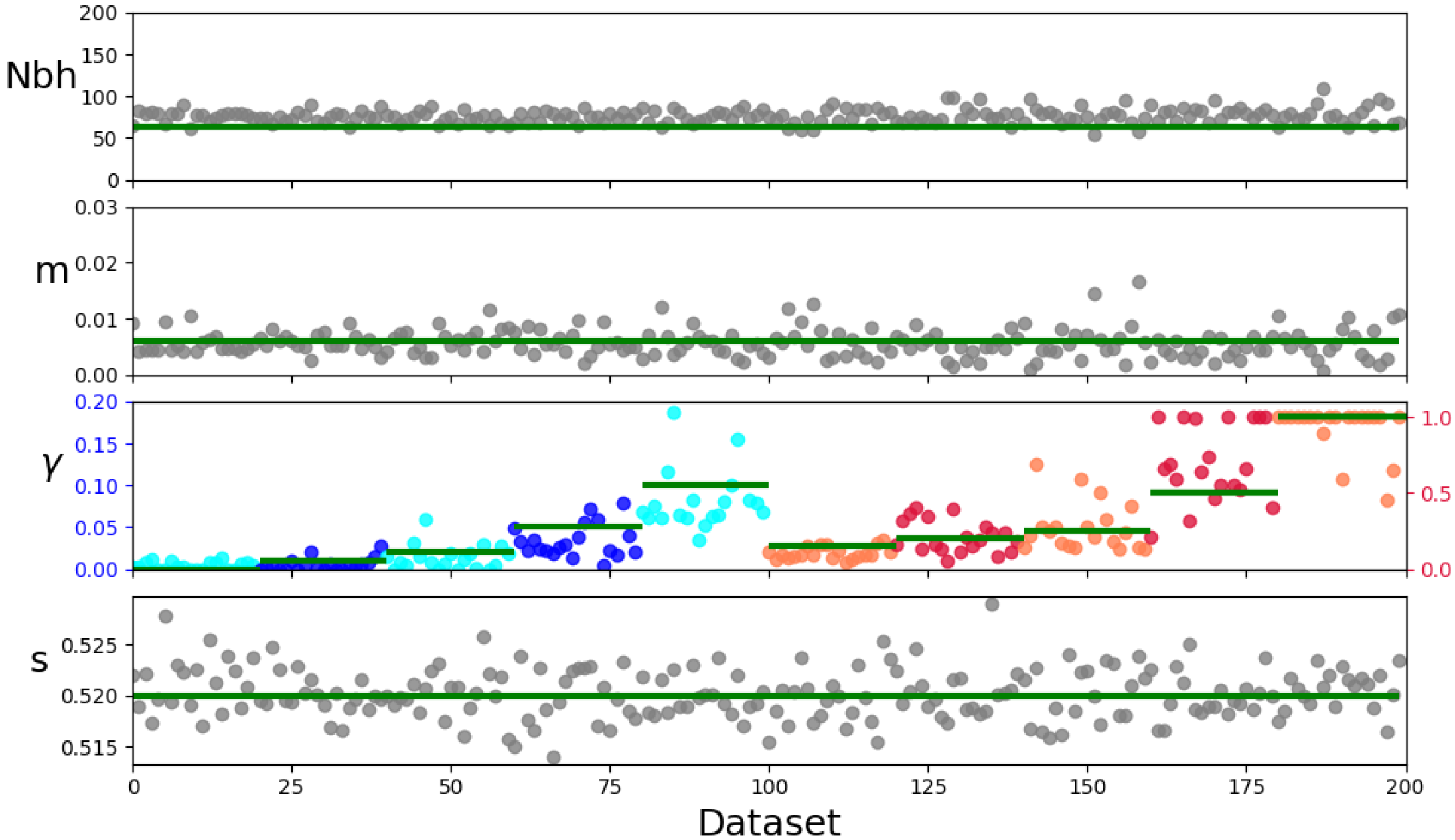
Parameter estimates on simulated data. For each replicate, we simulated a population of 60 × 40 individuals spaced one node apart, and then applied the inference scheme to fit neighborhood size Nbh, barrier strength *γ*, scaled long-distance migration rate *m*, as well as the allele frequency variance parameter *s*. To keep run-times manageable, we binned individuals into 2 × 2 demes. We simulated 10 different barrier strengths (*γ* = 0,0.01,0.02,0.05,0.1,0.15,0.2,0.25,0.5,1.0), with 20 replicate runs each. Every dot represents an estimate for one such replicate. Horizontal lines depict the true parameters used in the simulations (*σ* = 1, *D*_*e*_ = 5, *μ* = 0.003, *σ*(*p̄*) = 0.1). For better visibility we split up the barrier plot into two parts with different axes (blue and red).

**Figure 6.**
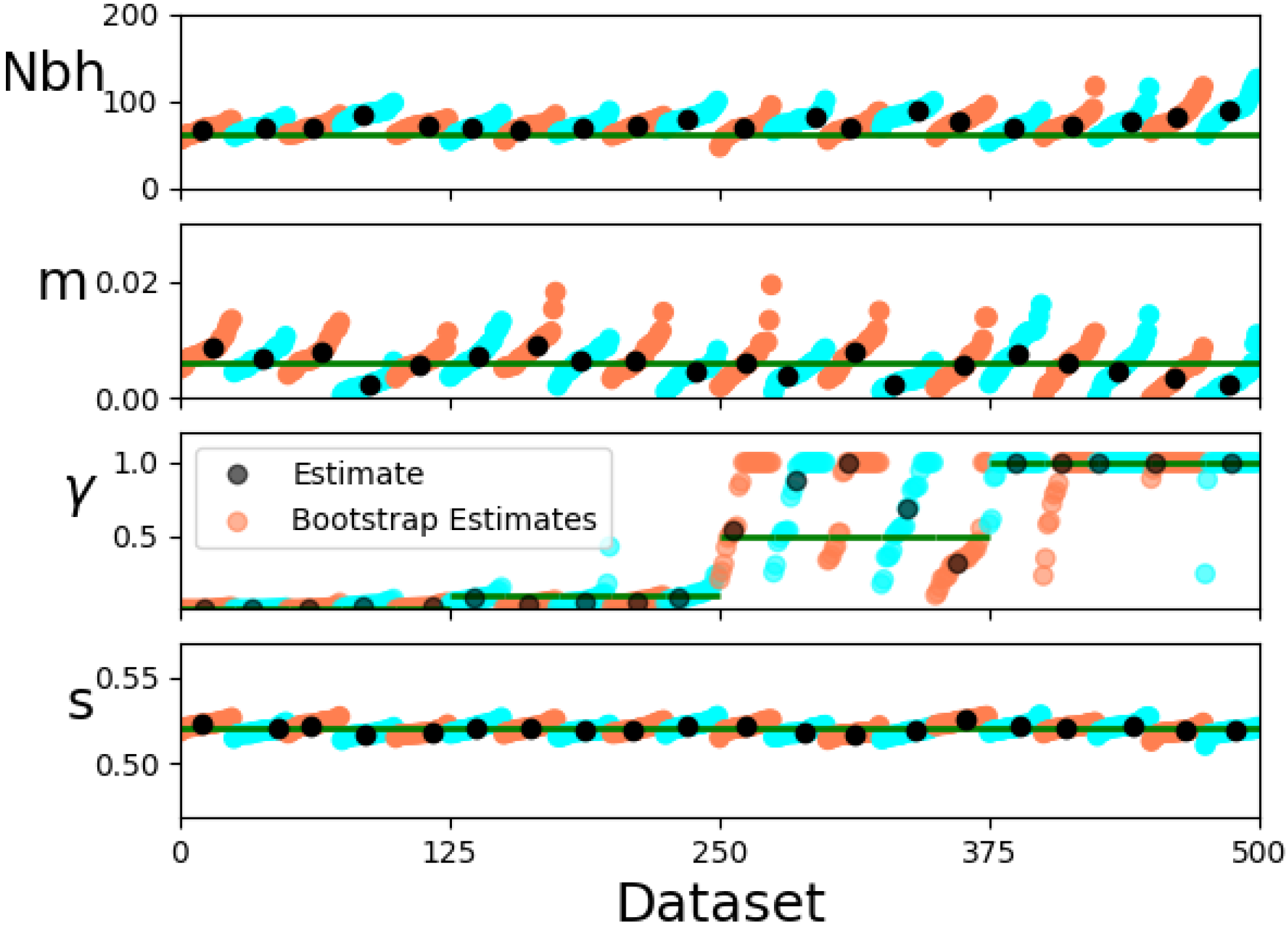
Bootstrap estimates on simulated data: We simulated 60 × 40 individuals on a grid spaced one distance unit apart. We simulated 5 replicates of 4 different barrier strengths (*γ* = 0,0.1,0.5,1.0). For each of the 20 data-sets we inferred the parameters and additionally did 20 estimates when bootstrapping over loci. Horizontal lines depict the true parameters used in the simulations (*σ* = 1, *D*_*e*_ = 5, *μ* = 0.003, *σ*(*p̄*) = 0.1)

**Figure 7.**
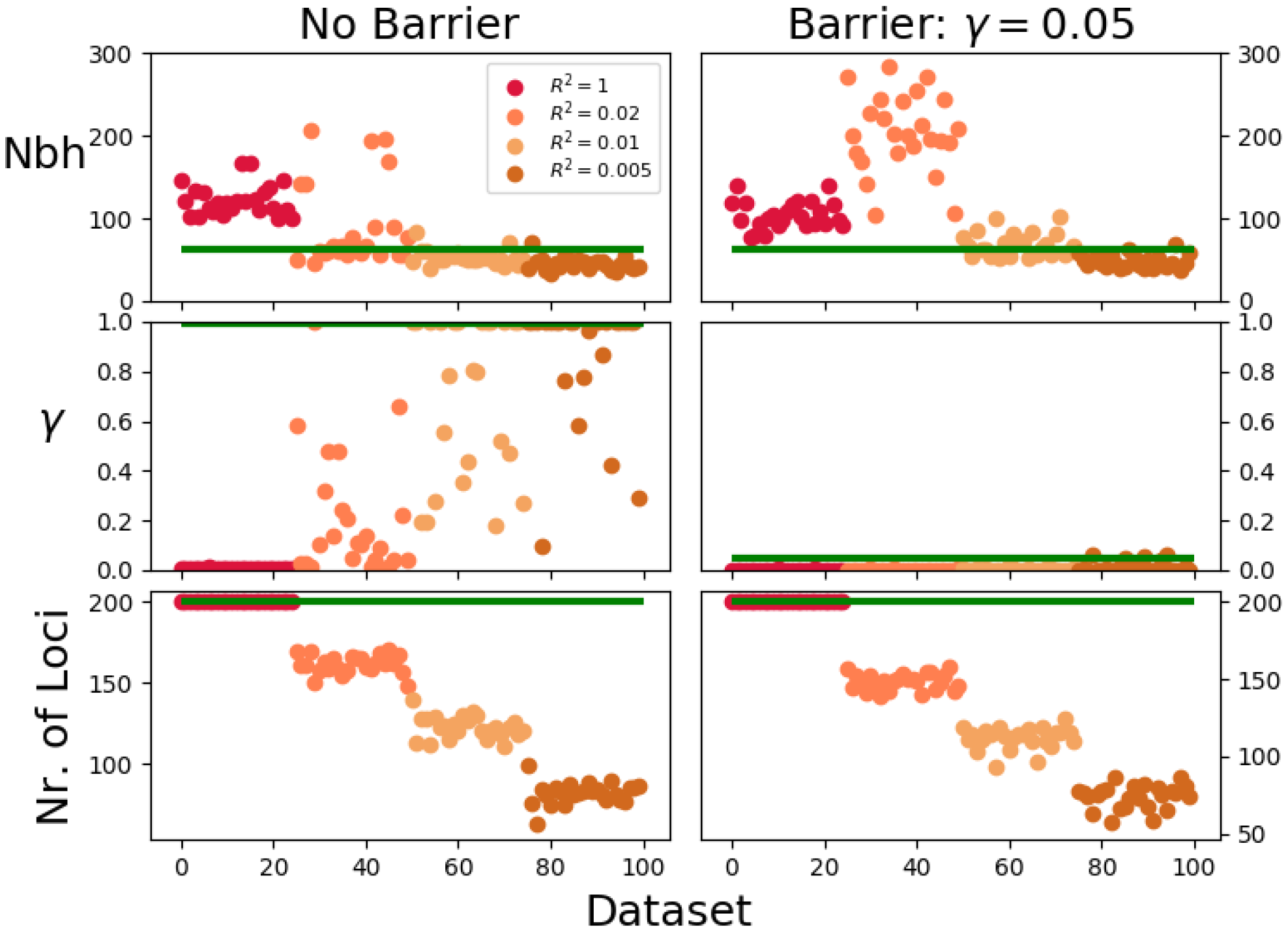
Parameter estimates in a simulation of secondary contact. The two ancestral populations allele frequencies were drawn independently from a Gaussian with standard deviation 0.1 around an overall mean of 0.5. We simulated a population of 50 × 20 individuals spaced two dispersal units apart, with a barrier in the middle of the *x*-axis, and secondary contact 100 generations ago. We simulated 25 replicates of two scenarios after contact. Left: No barrier to gene flow after contact. Right: A strong barrier to gene flow (*γ* = 0.05). We then filtered loci that were correlated with either *x*- or *y*-axis coordinates more than four different *R*^2^ values and redid inference for these replicates. The bottom row depicts the number of filtered loci that were left after filtering.

#### Unknown barrier locations

Our inference scheme assumes that the location of the putative barrier is known a priori. In practice one might not always have this information, or one perhaps wants to test the hypothesis of barriers in different locations. In this case, one can repeatedly apply the inference scheme and try out different potential barrier positions. When testing this approach on simulated data, the inference scheme only inferred a strong barrier near the true position (Fig. 8). The estimate uncertainties on the habitat edges are inflated. This effect is caused by limited power to infer barriers near sampling edges: One needs a sufficient number of samples on both sides of a barrier to estimate the strength of a barrier.

**Figure 8.**
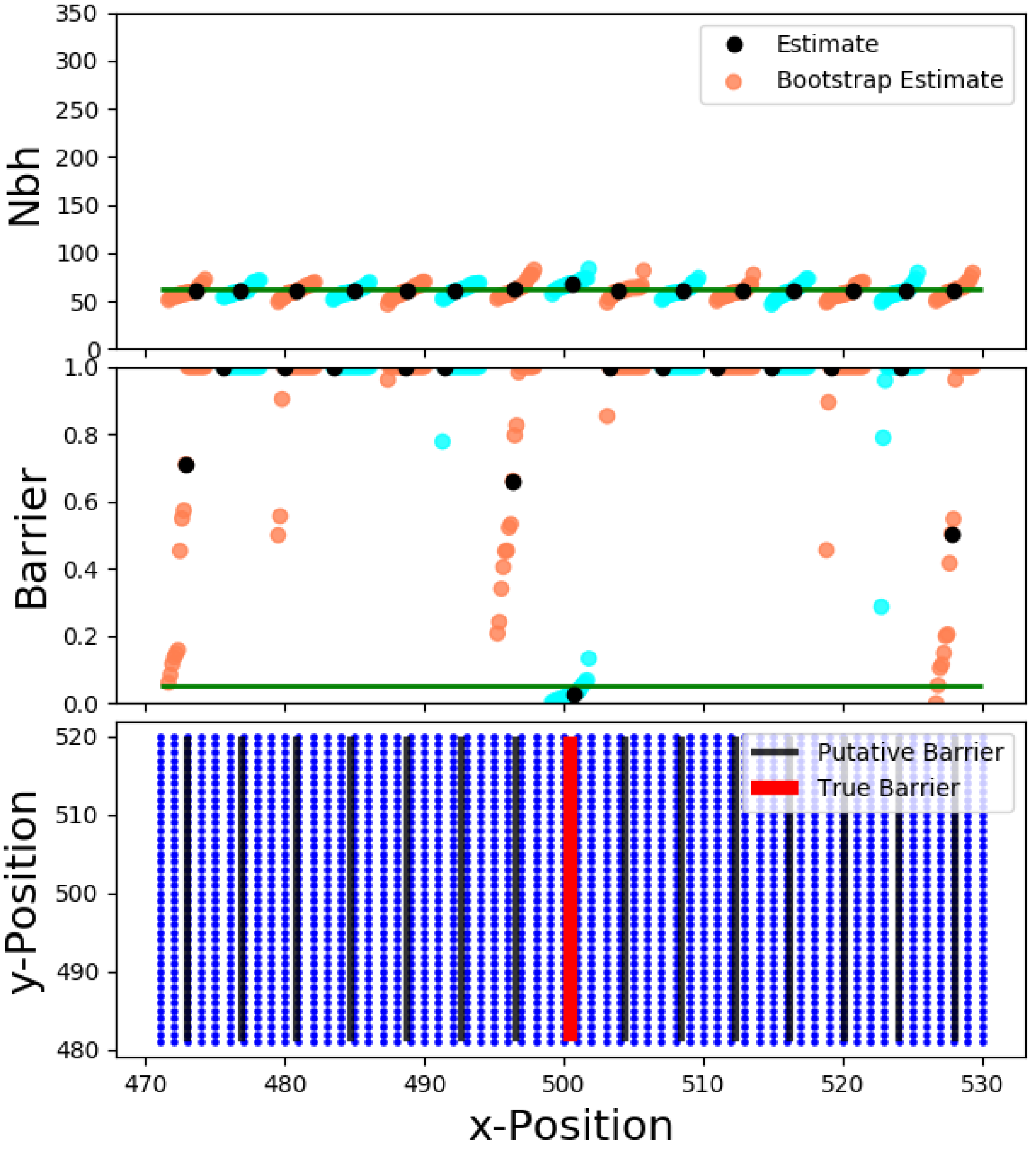
Testing various putative barrier positions on synthetic data. We simulated a single dataset as in Fig. 5 with a barrier strength of *γ* = 0.05. We applied our inference scheme to fit 15 putative barrier locations, and bootstrapped 20 times over loci for each putative barrier to visualize the uncertainty of the inferred parameters.

### Hybrid Zone Analysis

We observed a clear isolation by distance pattern across the *Antirrhinum* hybrid zone (Fig. 9). On average, mean identity by state for nearby plants is elevated by about two percent above the background level, and falls away with increasing pairwise distance, most rapidly over the first 2000 meters. Our inference scheme fits this pattern well, with an estimated neighborhood size of 188 (95% bootstrap confidence intervall: 120 – 240).

**Figure 9.**
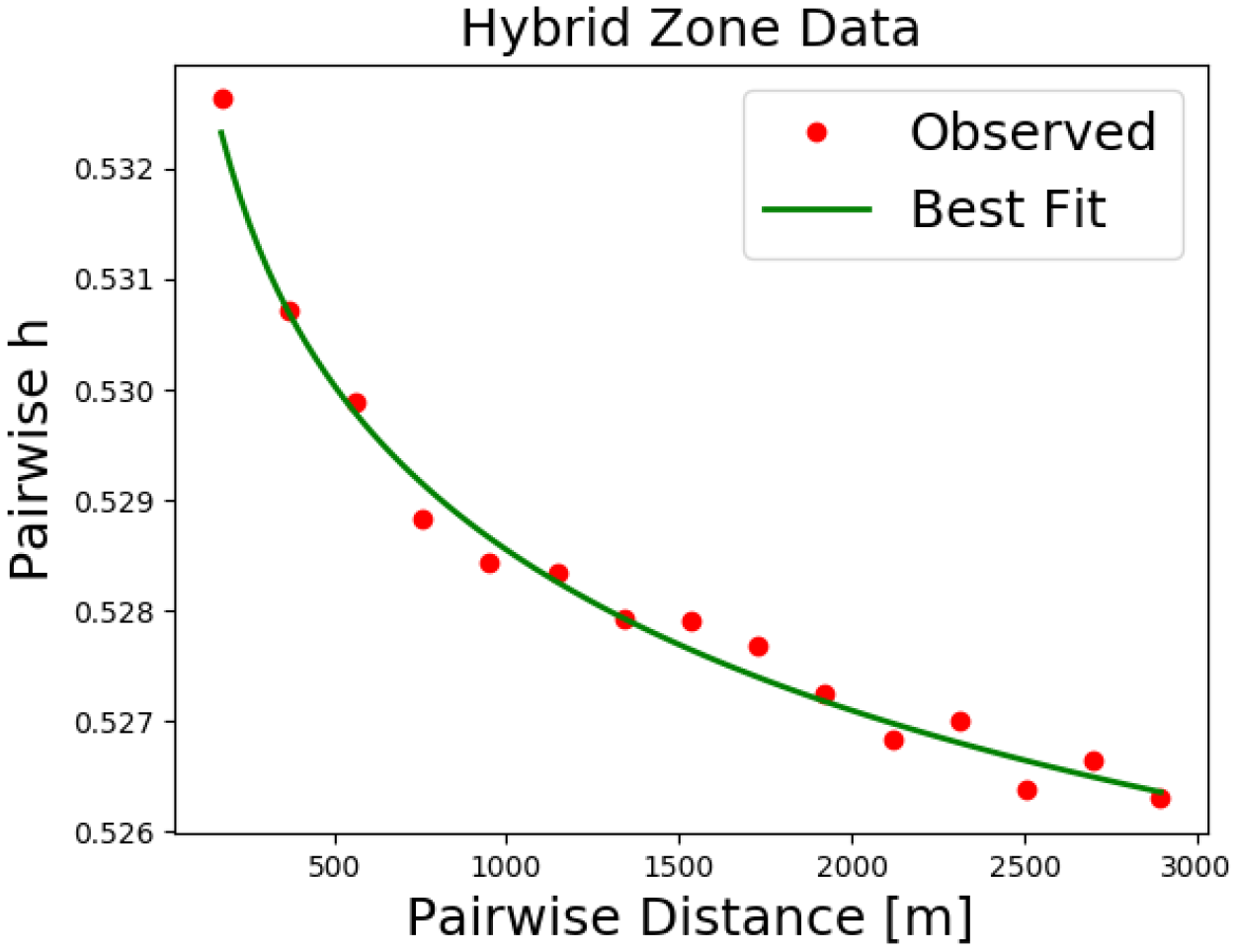
Decay of pairwise homozygosity with geographic distance for hybrid zone data. All pairwise homozygosities for the filtered dataset were binned according to pairwise distance. We also plot the best fit (Nbh = 188, m = 4.4 · 10^−4^,s = 0.5247). Formula 10 can be used to translate pairwise homozygosity into pairwise *F*.

For putative barriers in the center of the hybrid zone, our inference scheme estimates no barrier (*γ* > 1). Bootstrap estimates rarely fall below *γ* = 0.5, and all of them are above *γ* = 0.1. When testing for barriers towards the flank of the hybrid zone, estimates get more variable. This signal likely reflects the lower power to infer barriers, as most sampled plants are in the center of the hybrid zone (58.7% of the samples are from within 500 meter of the flower color transition).

Obviously there are demographic complications that are not captured by our model, such as heterogeneities in plant distributions and density. The population is not distributed uniformly in two dimensions, as plants are often found in patches of suitable habitat (Fig. 10). However, our analysis indicates that isolation by distance patterns are neither strongly influenced by the cardinal direction or relative plant positions, nor the geographic location within the hybrid zone (Supp. Text. 3). These observations imply that our model assumptions are not grossly violated.

**Figure 10.**
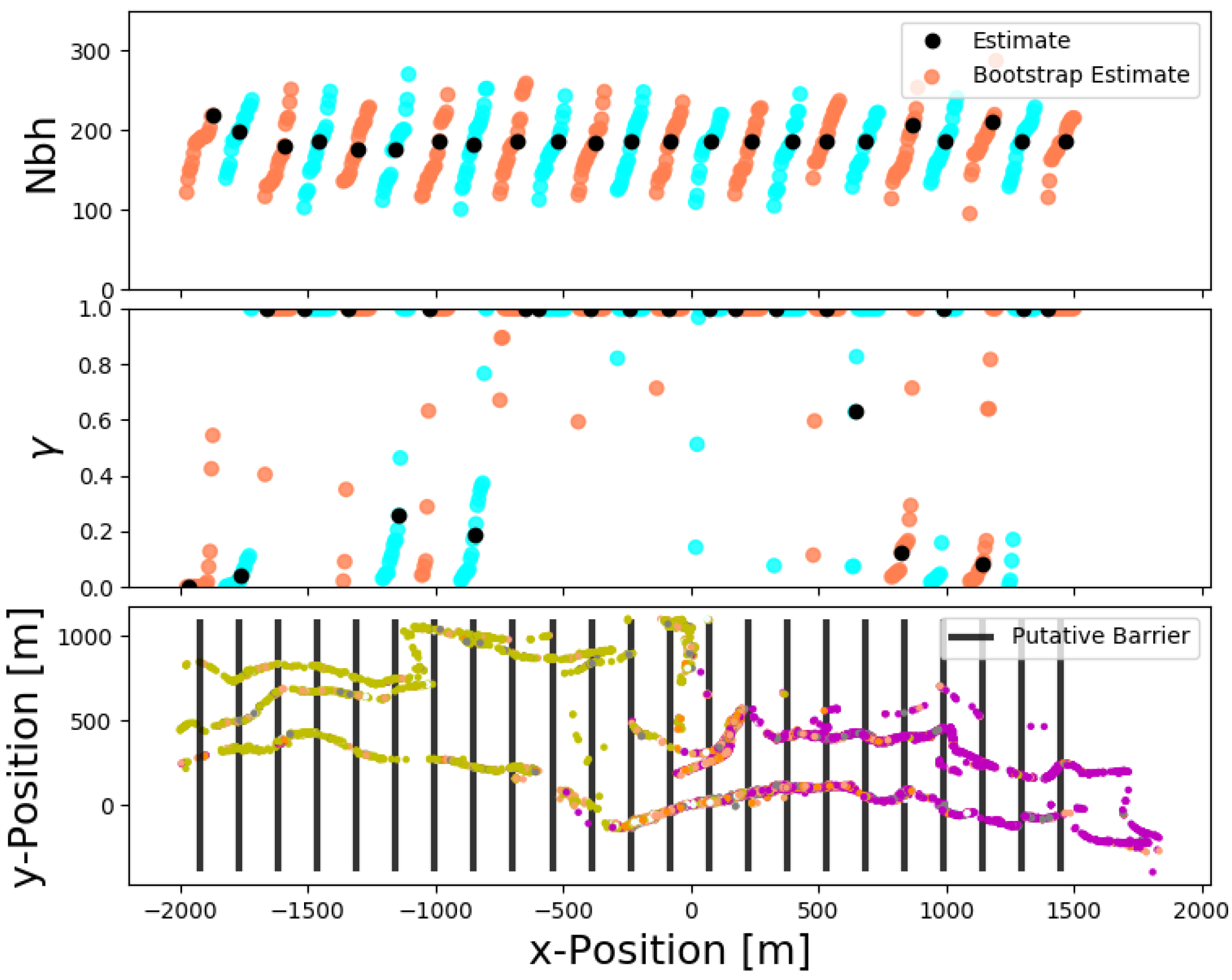
Inference of various putative barrier positions on data from a *Antirrhinum majus* population. We used our inference scheme to fit 15 barrier locations, and bootstrapped 20 times over loci for each putative barrier (bootstrap estimates are shown in orange and cyan). Bottom: Geographic distribution of 12389 samples. The color of each dot represents flower-color phenotype.

Given the overall good fit of isolation by distance with our inference scheme (Fig. 9), our results indicate that there is no strong genome wide barrier to contemporary gene flow that coincides with the flower color transition. As such a strong barrier would require many barrier loci spaced densely throughout the genome (Barton and Bengtsson 1986), this result comes as no surprise. At the moment, no other traits apart from flower color are known to be divergent across the hybrid zone, despite much work to detect them (Personal Communication: Maria Melo).

Previous results suggest the presence of a barrier to exchange of flower color alleles (Whibley *et al.* 2006) and indicate that selection maintains differences in flower color (Ellis 2016). Therefore, we also applied our method to a subset of polymorphic markers in the genetic neighborhood of two genes known to affect flower color variation in the hybrid zone, *Rosea* and *Eluta*. However, bootstraps estimates varied widely for all tested barrier locations (results not shown), which indicates that there is not enough signal in the data. This is likely due to the low number of suitable SNP markers without steep allele frequency clines near this region in our dataset (< 10). Simulations confirmed that for this low number of markers there is not enough power to detect even strong barriers (Supp. Text 1).

## Discussion

To our knowledge, our scheme is the first method that infers the strength of the barrier based on an explicit spatial population genetic model. There are several similarities with the inference method BEDASSLE (Bradburd *et al.* 2013), which aims to disentangle the effects of geographical isolation by distance and differences in ecological variables. This scheme is not based on an explicit population genetic model, but rather fits the decay of genetic similarity with distance to a heuristic formula. In light of possibly very complex demographic structure this method is not necessarily worse than fitting our spatial model. However, this approach does not take the increased covariances on the same side of a barrier into account, and does not make use of some valuable signal because of that. Moreover, it uses a MCMC approach that is based on a model of Gaussian random Fields, which is computationally too expensive to apply to hundreds or even thousands of individuals or demes. In contrast, our method is well suited to data sets of this magnitude.

Another widely used method to infer barriers to gene flow, Geneland, clusters individuals using their explicit geographic coordinates (Guillot *et al.* 2005). Safner *et al.* (2011) identified it as one of the most potent methods to infer barriers to gene flow. Therefore, we tested its performance on some of the datasets which we have generated to test our scheme (Supp. Text 2). As described previously (Guillot and Santos 2009; Safner *et al.* 2011), Geneland’s ability to accurately infer barriers to gene flow decreases when isolation by distance patterns are present, as its underlying model assumptions of discrete populations with well-defined allele frequencies are violated. Indeed, it fails to detect a barrier to gene flow in our test data sets, that all exhibit such isolation by distance patterns (Supp. Text 2). In contrast, the method introduced here can give accurate estimates of the barrier strength in these scenarios. It is not confounded by isolation by distance patterns; it in fact relies on the presence of this signal. Our method can therefore be seen as complementary to Geneland.

In contrast to BEDASSLE, Geneland and most other existing methods, which all heuristically describe the strength of the parameter, the inference scheme introduced here fits an explicit spatial population genetics model. It corresponds to Nagylaki’s *γ* (Nagylaki 1976), whose inverse 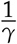 is equal to Barton’s barrier strength *B* (Barton and Bengtsson 1986). This correspondence makes the inferred barrier strength *γ* a parameter that is directly interpretable in terms of population genetic theory. For instance, the parameter 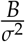, which has dimension of time, must be large to retard the spread of even a neutral allele (Barton and Bengtsson 1986).

Our method can reliably estimate the strength of strong barrier (*γ* ⪅ 0.1), but there is little power to distinguish between a weak (*γ* ⪆ 0.25) and no barrier (Fig. 5 and Fig. 6). The underlying reason is not a shortcoming of our inference scheme, but the fact that relatively weak barriers do not significantly affect the spread of ancestry (Fig. 2). Therefore, it is infeasible to estimate the strength of weak barriers to gene flow, simply because they do not have a significant effect on allele frequency covariances. This effect was already observed by Barton *et al.* (2013), who found that in two spatial dimensions the effect of a barrier starts to have appreciable effects on the spatial pattern of genetic marker alleles when barrier strength *B* (which corresponds to 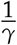) is ≈ *σ*.

## Outlook

The inference scheme introduced here fits a linear barrier, the most straight-forward model for a barrier in two dimensions. We used an analytical formula (Eq. 5) to model the spread of ancestry, which in turn allows one to reduce calculations for pairwise *F* to a single numerical integral (Eq. 7). However, in practice barriers might be geographically more complex. Most more complicated scenarios will not allow for a simple formula, and calculations for the chance of recent co-ancestry become more challenging. The techniques outlined here can still be applied, but ancestry has to be traced back with explicit simulations. This numerical problem seems to be within reach of present day computational power, and would be a salient extension to our model.

Our method fits allele frequencies, which can be confounded by deeper ancestral patterns. By filtering loci that show large scale geographic variation, one can in principal remove some of this ancestral genetic structure, but by doing so one might accidentally remove true signal as well. This problem can be a severly confounding factor when applying the inference method to scenarios were ancestral structure is present, for instance zones of putative secondary contact.

One promising way to overcome this problem would be to base inference on identity by descent blocks, the direct genetic traces of recent co-ancestry (Browning and Browning 2012). As blocks of ancestral genetic material are split up at a constant rate by recombination, the probability of sharing a block of length *l* decays exponentially back in time (Ralph and Coop 2013). Therefore, blocks longer than 5 cM, say, are very unlikely to originate from co-ancestry older than 100 generations, even under relatively extreme demographic scenarios. Moreover, the length of the blocks contains information about the time of coalescent. Identifying such blocks is a non-trivial task, in particular when only un-phased genotype data is available (Browning and Browning 2012). It requires dense genotype data and linkage information. But in cases where IBD blocks can be robustly called - as already possible for humans and some model organisms - an inference scheme based on this signal holds great potential. Our method to model the spread of ancestry can be combined with formulas for block sharing (Ralph and Coop 2013; Ringbauer *et al.* 2017) to calculate the expected number of shared IBD blocks in presence of a barrier. These results could be used to fit observed block sharing data.

Summarizing, our method is only a first step to robustly infer barriers to gene flow from genotype data. The techniques outlined here can be expanded in various directions to better deal with the complexities of real data, and to make full use of opportunities within the era of population genomics. We hope that this will ultimately lead to a better understanding of barriers to gene flow within many natural populations.

## APPENDIX

Here we give the full formula we fit, and describe the rescaling to a set of independent effective parameters. Let *x, y* denote the *x*-coordinate of the samples, and Δ*y* their separation along the y-axis. For the identity by state on different sides of the barrier plugging into formula Eq. 7 gives for *F*(*x*, *y*, Δ*y*):

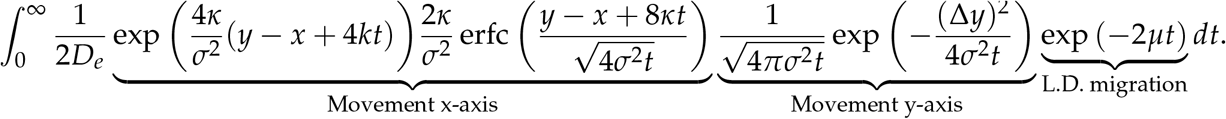

We now rescale time, such that t′ = *σ*^2^*t*. The integral (*dt*′ = *σ*^2^*dt*) transforms to:

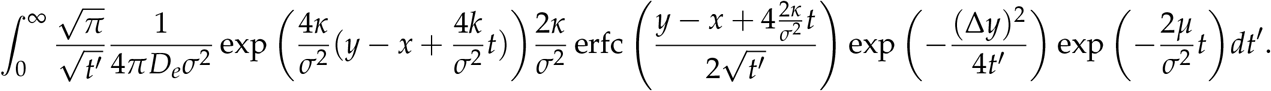

Defining Nbh:= 4*πD*_*e*_*σ*^2^, 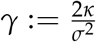 and 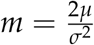 gives the full formula used for inference:

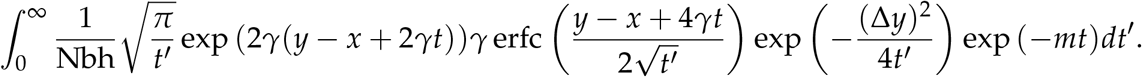

The formula for same sides of the barrier is rescaled analogously. The additional term in the integrand becomes:

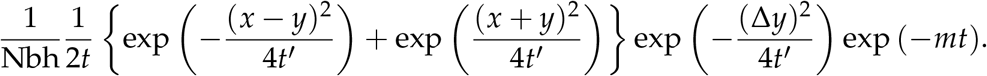

## Literature Cited

Aguillon, S. M., J. W. Fitzpatrick, R. Bowman, S. J. Schoech, A. G. Clark, G. Coop, and N. Chen, 2017 Deconstructing isolation-by-distance: the genomic consequences of limited dispersal. PLoS Genetics 13: e1006911.

Barton, N., 1979 Gene flow past a cline. Heredity 43: 333–339.

Barton, N., 2008 The effect of a barrier to gene flow on patterns of geographic variation. Genetics research 90: 139–149.

Barton, N. and B. O. Bengtsson, 1986 The barrier to genetic exchange between hybridising populations. Heredity 57: 357–376.

Barton, N., A. Etheridge, J. Kelleher, and A. Véber, 2013 Inference in two dimensions: allele frequencies versus lengths of shared sequence blocks. Theoretical population biology 87: 105–119.

Barton, N. H., F. Depaulis, and A. M. Etheridge, 2002 Neutral evolution in spatially continuous populations. Theoretical population biology 61: 31–48.

Barton, N. H. and G. M. Hewitt, 1985 Analysis of hybrid zones. Annual review of Ecology and Systematics 16: 113–148.

Bradburd, G. S., P. L. Ralph, and G. M. Coop, 2013 Disentangling the effects of geographic and ecological isolation on genetic differentiation. Evolution 67: 3258–3273.

Browning, S. R. and B. L. Browning, 2012 Identity by descent between distant relatives: detection and applications. Annual review of genetics 46: 617–633.

Cercueil, A., O. François, and S. Manel, 2007 The genetical bandwidth mapping: a spatial and graphical representation of population genetic structure based on the wombling method. Theoretical population biology 71: 332–341.

Dupanloup, I., S. Schneider, and L. Excoffier, 2002 A simulated annealing approach to define the genetic structure of populations. Molecular Ecology 11: 2571–2581.

Ellis, T. J., 2016 The role of pollinator-mediated selection in the maintenance of a flower color polymorphism in an Antirrhinum majus hybrid zone. Ph.D. thesis, IST Austria.

Falush, D., M. Stephens, and J. K. Pritchard, 2003 Inference of population structure using multilocus genotype data: linked loci and correlated allele frequencies. Genetics 164: 1567–1587.

Fisher, R. A., 1937 The wave of advance of advantageous genes. Annals of eugenics 7: 355–369.

Grebenkov, D. S., D. Van Nguyen, and J.-R. Li, 2014 Exploring diffusion across permeable barriers at high gradients. i. narrow pulse approximation. Journal of Magnetic Resonance 248: 153–163.

Guillot, G., A. Estoup, F. Mortier, and J. F. Cosson, 2005 A spatial statistical model for landscape genetics. Genetics 170: 1261–1280.

Guillot, G., R. Leblois, A. Coulon, and A. C. Frantz, 2009 Statistical methods in spatial genetics. Molecular Ecology 18: 4734–4756.

Guillot, G. and F. Santos, 2009 A computer program to simulate multilocus genotype data with spatially autocorrelated allele frequencies. Molecular Ecology Resources 9: 1112–1120.

Hardy, O. J. and X. Vekemans, 1999 Isolation by distance in a continuous population: reconciliation between spatial autocorrelation analysis and population genetics models. Heredity 83: 145–154.

Malécot, G., 1948 mathematiques de l’heredite.

Manni, F., E. Guerard, and E. Heyer, 2004 Geographic patterns of (genetic, morphologic, linguistic) variation: how barriers can be detected by using monmonier’s algorithm. Human biology 76: 173–190.

Meirmans, P. G., 2012 The trouble with isolation by distance. Molecular ecology 21: 2839–2846.

Nagylaki, T., 1976 Clines with variable migration. Genetics 83: 867–886.

Nagylaki, T., 1978 A diffusion model for geographically structured populations. Journal of Mathematical Biology 6: 375–382.

Nagylaki, T., 1988 The influence of spatial inhomogeneities on neutral models of geographical variation: I. formulation. Theoretical Population Biology 33: 291–310.

Ralph, P. and G. Coop, 2013 The geography of recent genetic ancestry across europe. PLoS biology 11: e1001555.

Ringbauer, H., G. Coop, and N. H. Barton, 2017 Inferring recent demography from isolation by distance of long shared sequence blocks. Genetics 205: 1335–1351.

Rousset, F., 1997 Genetic differentiation and estimation of gene flow from f-statistics under isolation by distance. Genetics 145: 1219–1228.

Rousset, F., 2002 Inbreeding and relatedness coefficients: what do they measure? Heredity 88: 371–380.

Safner, T., M. P. Miller, B. H. McRae, M.-J. Fortin, and S. Manel, 2011 Comparison of bayesian clustering and edge detection methods for inferring boundaries in landscape genetics. International Journal of Molecular Sciences 12: 865–889.

Slatkin, M., 1993 Isolation by distance in equilibrium and non-equilibrium populations. Evolution 47: 264–279.

Whibley, A. C., N. B. Langlade, C. Andalo, A. I. Hanna, A. Bangham, C. Thébaud, and E. Coen, 2006 Evolutionary paths underlying flower color variation in antirrhinum. Science 313: 963–966.

Wilkins, J. F., 2004 A separation-of-timescales approach to the coalescent in a continuous population. Genetics 168: 2227–2244.

Womble, W. H., 1951 Differential systematics. Science 114: 315–322.

Wright, S., 1943 Isolation by distance. Genetics 28: 114.

